# Temporal and spatial asymmetries during stationary cycling cause different feedforward and feedback modifications in the muscular control of the lower limbs

**DOI:** 10.1101/370262

**Authors:** Magdalena Zych, Ian Rankin, Donal Holland, Giacomo Severini

**Author notes:** Correspondence: Giacomo Severini, PhD School of Electrical and Electronics Engineering, University College Dublin, Belfield, Dublin 4, Ireland.

## Abstract

Motor adaptations are useful for studying the way in which the lower limbs are controlled by the brain. However, motor adaptation paradigms for the lower limbs are typically based on locomotion tasks, where the necessity of maintaining postural stability is the main driver of adaptation and could possibly mask other underlying processes. In this study we investigated whether small temporal or spatial asymmetries can trigger motor adaptations during stationary cycling, where stability is not directly compromised. Fourteen healthy individuals participated in two experiments: in one of the experiments the angle between the crank arms of the pedals was altered by 10° to induce a temporal asymmetry; in the other the length of the right pedal was shortened by 2.4 cm to induce a spatial asymmetry. We recorded the acceleration of the crank arms and the EMG signals of 16 muscles (8 per leg). The analysis of the accelerometer data was used to investigate the presence of motor adaptations. Muscle synergy analysis was performed on each side to quantify changes in neuromuscular control. We found that feedforward motor adaptations are present in response to temporal asymmetries and are obtained by progressively shifting the activation patterns of two synergies on the right leg. Spatial asymmetries appear to trigger a feedback-driven response that does not present an aftereffect and is not consistent with a motor adaptation. This response is characterized by a step-like decrease in activity in the right gastrocnemius when the asymmetry is present and likely reflects the altered task demands.

**New and Noteworthy:** The processes driving lower limb motor adaptations are not fully clear, and previous research appears to indicate that adaptations are mainly driven by stability. Here we show that lower limb adaptations can be obtained also in the absence of an explicit balance threat. We also show that adaptations are present also when kinematic error cannot be compensated for, suggesting the presence of intrinsic error measures regulating the timing of activation of the two legs.

## Introduction

When performing repetitive movements such as walking and cycling, the human central nervous system coordinates different neural commands to form patterns of alternating muscular activations that are mostly automated. The planning and voluntary adjustments of the motor plan take place in supraspinal areas and the cortex (Caggiano et al. 2018; Drew 1988; Jordan et al. 2008; Takakusaki 2013) while the muscular pattern formation and the timing of activation is mostly regulated at the spinal level (Caggiano et al. 2018; Grillner 2003; Kiehn 2016; 2006). During walking, if a visual discrepancy (Logan et al. 2014), a sudden obstacle (Schillings et al. 2000) or a continuous perturbation (Cajigas et al. 2017) disrupts the task in a relevant way, these automated patterns are modulated to overcome or reject the disturbance. Fast interferences (e.g. tripping into an obstacle or slipping on a slippery surface) are compensated through spinal reflex responses (McDonagh and Duncan 2002; Schillings et al. 2000; van der Linden et al. 2007), while continuous perturbations or modifications of the movement environment require for the motor plan to be progressively updated to reject the error in a phenomenon often referred to as locomotor adaptation (Torres-Oviedo et al. 2011).

While our knowledge of spinal reflexes and their role during locomotion is substantial (Kandel et al. 2000; Pearson 2004; Rossignol et al. 2006), the processes behind the generation of locomotor adaptations are not fully understood yet. Most of our knowledge on lower limb and locomotor adaptations derives from studies based on actual walking disturbances administered either by using robotic systems (Cajigas et al. 2017; Emken et al. 2007; Emken and Reinkensmeyer 2005) or by altering the walking environment, such as in the split-belt treadmill paradigm (Choi and Bastian 2007; Prokop et al. 1995; Reisman et al. 2005). These studies have shown that motor adaptations during locomotion are generated mostly to maintain postural stability (Cajigas et al. 2017; Lam et al. 2006; Prokop et al. 1995), to reduce energy consumption (Emken et al. 2007; Finley et al. 2013) and due to a natural bias towards symmetry (Reisman et al. 2005). A recent work has shown that stability has primacy over the other drivers of adaptation (Cajigas et al. 2017), whereas perturbations altering postural stability are compensated even if they lead to an increased energy consumption while perturbations that do not alter postural stability are ignored in favour of a more economical gait pattern. Nevertheless, it is still not fully clear which circuits are adapted during locomotor adaptations and, since most experimental paradigms alter the balance requirements during the task, if such adaptations are hardwired or a response to a behavioral trigger such as maintaining a long-term stable gait pattern.

This latter point is specifically of interest given the fact that locomotor adaptations have proved to be a valuable tool for clarifying the organization of the neural circuits controlling the lower limbs during repetitive movements in humans. In their seminal work, Choi and Bastian (Choi and Bastian 2007) used the split-belt treadmill paradigm to study the organization of the hypothetical Central Pattern Generator (CPG) circuits in the human spinal cord. Such CPG circuits have been proposed as constituted by spatial components coordinating the activity of the different muscles together and rhythmic components coordinating the timing of activation of the spatial components (Kiehn 2006; McCrea and Rybak 2008; Zhong et al. 2012). Several studies have proposed that muscle synergies extracted by applying factorization algorithms to datasets constituted by large numbers of EMG signals may functionally model the two-layer structure of the CPGs in humans (Dominici et al. 2011; Ivanenko et al. 2005; Lacquaniti et al. 2012; MacLellan et al. 2014).

In this work we propose a novel method for the study of lower limb motor adaptations based on asymmetric cycling. In this method we introduce small temporal or spatial asymmetries in the cycling pattern of healthy individuals during exercises on a stationary bike to verify the presence of locomotor adaptations and study the associated neuromuscular correlates (MacLellan et al. 2014). Temporal asymmetries are introduced by slightly altering the relative angle between the crank arms of the cycling system, while spatial asymmetries are introduced by altering the length of one of the two crank arms. This approach allows us to study automatic reactions, if any occur, to alterations in the normal temporal and spatial organization of repetitive lower limb movements without introducing perturbations that may directly affect balance. Conceivably, introducing an asymmetry in the timing of activation of the two legs disrupts the coordination of the rhythmic components of the CPGs. On the other hand, the spatial asymmetry modifies the muscular requirements for the task and may translate in modifications in the layer of the CPGs regulating the co-recruitment of the different muscles. Observing motor adaptations in one or both of these components may help to identify which of the neural circuits that control lower limb movements are automatically adapted even in the absence of a behavioral primer such as a balance threat. Herein we studied the presence of motor adaptation during the previously described two asymmetric cycling tasks by tracking changes in acceleration patterns in the crank arm and in the temporal and spatial components of the muscle synergies (d’Avella et al. 2003) associated with both experiments. Muscle synergy analysis was performed on the muscles of each leg separately to get an insight in the activity of the different components of the unilateral CPGs controlling the lower limbs.

## Materials and Methods

### Participants

Fourteen healthy individuals (7 females; 26.1 ± 3.0 years old, 171.6 ± 6.7 cm, 64.2 ± 7.3 kg) volunteered to participate in the experiments. All participants reported their right leg to be the dominant one, intended as the leg they would use to kick a ball. Inclusion criteria consisted of the absence of neurological, orthopaedic or cognitive impairments that would in any way affect the execution of the experiment. Only two out of fourteen subjects routinely used clipless pedals when cycling (recreationally or for commute), while all the others utilized standard pedals. All the experimental procedures described in the following have been approved by the Ethical Committee of University College Dublin and have been conducted according to the WMA’s declaration of Helsinki. All subjects gave written informed consent before participating in this study.

### Experimental Procedures

All tests were performed by each participant in a single experimental session. Participants were instructed to refrain from vigorous physical activity within 2 hours before testing. During the experimental session each participant underwent two sessions of cycling, each relative to a different symmetry perturbation. All cycling exercises were performed using a standard training cycloergometer (Lode BV, Groningen, The Netherlands). The pedals of the cycloergometer were equipped with straps that were used to solidly attach the feet of the subjects to the pedal during the experiments.

One symmetry perturbation (Angle experiment) consisted in a 10° clockwise (with respect to the right leg) modification in the angle between the two crank arms. This was achieved by using a custom metal plate directly attached to the end of the right crank arm (**Figure 1**). The other symmetry perturbation (Length experiment) consisted in a shortening of the right crank arm by 2.3 cm. This was obtained by attaching a commercial shortening system (Pulse Crank Shortener Highpath Engineering Ltd., Tring, UK) to the right crank arm (**Figure 1**). The “unperturbed” (where right is the perturbed side and left the unperturbed one) crank arm length was equal to 17 cm. Both asymmetries were designed to be very small and barely acknowledgeable by the subjects, in order not to modify too much the normal biomechanics of cycling and so that potential adaptation behaviors would be involuntary. In both experiments the participants were asked to cycle for 5 minutes without asymmetries (baseline, BL), followed by 10 minutes of asymmetric cycling (adaptation, AD) and concluded each experiment with 5 minutes of symmetric cycling (post-adaptation, PA, **Figure 1**). Between each phase of each experiment the participants were allowed 5 minutes of rest, that were used to modify the pedal setup (e.g. to add or remove the asymmetries). Participants were specifically asked not to walk during the resting period between AD and PA to avoid possible washouts. Between the two experiments each participant was given a 20 minutes break. The two experiments were randomized across subjects with half of the subjects performing the Angle experiment first. In all phases of both experiments the cycloergometer was set at a constant torque equal to 6.5 Nm. This value was chosen so to keep the participant engaged while minimizing the possibility of muscle fatigue. Subjects were asked to reach as soon as possible and maintain a recreational pace of 75 revolutions per minute (rpm) during all phases of the experiments. Visual feedback on their pace was presented to them on the digital screen of the cycloergometer. Before the beginning of the first experiment the position of the saddle was adjusted to 109% of the inseam length between saddle and pedal in a pushed down position. This distance ensures optimal performance and safety for the locomotor system (Hamley and Thomas 1967; Peveler et al. 2007). Throughout the duration of both experiments, subjects were asked to keep their arms on the handles of the cycloergometers and to limit movements of the trunk.

**Fig 1.**
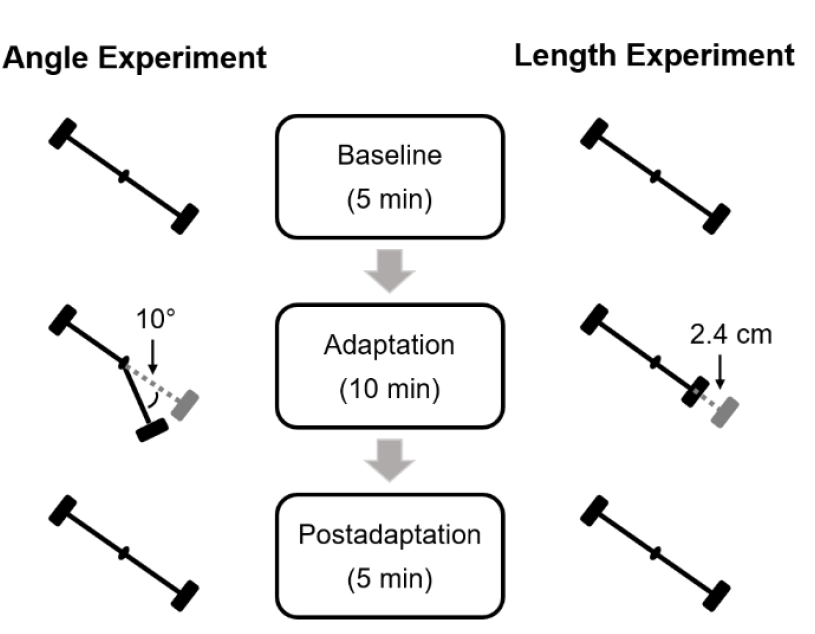
Experimental setup. The figure shows a representation of the lateral view of the right crank arm and pedal during the different phases of the two experiments. The asymmetry in the Angle experiment consisted of a 10 degrees clockwise (with respect to the right side) deviation in the relative angle between the crank arms of the cycloergometer. The asymmetry in the Length experiment consisted in a 2.4 cm shortening of the right crank arm. Both experiments consisted of 5 minutes symmetric cycling (baseline), followed by 10 minutes of asymmetric cycling (adaptation) and 5 minutes of symmetric cycling (postadaptation).

### Data recoding

The activity of 16 muscles (8 per leg) was recorded during both experiments using surface electromyography (sEMG). The electrode placement was performed at the beginning of the experimental session and data quality was checked through the different phases of both experiments. Wireless EMG sensors (Trigno Flex, Delsys, Boston, MA) were placed bilaterally (after the skin was prepared using alcohol), according to the SENIAM recommendations (Hermens et al. 1999), on the rectus femoris (RF), vastus lateralis (VL), tensor fasciae latae (TFL), gluteus maximus (GMax), biceps femoris (BF), tibialis anterior (TA), soleus (Sol) and gastrocnemius medialis (GM). The sampling frequency for the EMG sensors was set to 2000 Hz. One accelerometer (Trigno Flex sensors, Delsys, Boston, MA) was also placed on the left crank arm and sampled at 148 Hz. The accelerometer was placed, for both experiments, at half the length of the crank arm, so that it did not interfere with the metal attachments used to introduce the asymmetries.

A custom 6 cm flexible bend sensor was mounted under the left pedal in a bottom down position, so that the top 2 cm would bend each time the pedal was in its lowermost position. The data from the bend sensor were also sampled at 148 Hz and were used to determine the beginning of each cycle. All the sensors (EMG, accelerometers and bend sensors) where acquired synchronously using a custom software developed in Labview (Labview 2012, National Instruments, Austin, TX).

### Analysis of accelerometer data

All analysis routines were performed offline in MATLAB (MathWorks, Natick, MA). Biomechanical changes due to the introduction of the pedal asymmetries were estimated from the data recorded from accelerometer unit attached to the left crank arm. Changes in the duration (in time) of each cycle were also calculated from the data recorded from the flexible bend sensor. The *x* and *y* components (where *x* indicates the anteroposterior and *y* indicates the vertical direction) of the accelerometer data were segmented into individual cycles, identified as the interval between consecutive spikes in the flex sensor indicating the left pedal reaching the dead bottom of its trajectory, and length normalized to 100 points. The magnitude of the resultant acceleration on the sagittal plane vector was then computed for each segment as the square root of the sum of the squares of the *x* and *y* components of acceleration. As both the speed and the torque of each cycling exercise were fixed, changes in the accelerometer data were investigated as changes in the shape of the sagittal acceleration pattern across the different cycles. Specifically, we used as metric the position of the minimum sagittal acceleration that was determined to be robust in capturing asymmetry-driven changes in preliminary analyses (data on other metrics not shown). To investigate for possible exponential adaptive behaviors, as the one usually expected during locomotor adaptation and washout of adaptation (Cajigas et al. 2017), an exponential function was fitted on the median (across subjects) data from the AD and PA phases. The exponential function was fitted using a least square method and was in the form:

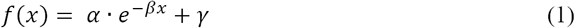

The time constant of the exponential, intended as the number of cycles needed for the exponential to reach its plateau, was estimated as 3/*β*. We also analysed for differences in the position of the peak acceleration during different phases of the experiments, nominally late BL, early AD, late AD and early PA. In this and all subsequent analyses “early” sections were constituted by the average of the first 50 cycles while “late” sections by the average of the last 50 cycles of a specific phase of each experiment. Changes in the average position of the peak acceleration were evaluated, across the different sections of both experiments, using Friedman’s test and a post-hoc analysis based on Dunn-Sidak’s criterion (non-normality of the data was assessed using the Kolmogorov-Smirnov test).

### EMG data processing and muscle synergies estimation

The recorded EMG signals were bandpass filtered (30-450 Hz) with a 4^th^-order band-pass Butterworth filter and rectified. The EMG envelopes were obtained from the data by filtering with a 3^rd^-order low-pass Butterworth filter with cut-off frequency of 5 Hz (De Marchis et al. 2015). The envelopes were then segmented into individual cycles using the same procedure utilized for segmenting the accelerometer data. Cycles with artefacts or abnormal, non-physiological, spiking activity were removed from subsequent analyses. All cycles of each channel were then amplitude normalized by dividing them by the average maximal value of the envelopes calculated from all the cycles from the baseline phase of each experiment (De Marchis et al. 2013; De Marchis et al. 2015). All the segmented cycles were then length-normalized by interpolating them to 100 points over a cycle. Muscle synergies were then extracted using the Non-Negative Matrix Factorization (NNMF) algorithm (Lee and Seung 2001). The algorithm decomposes the matrix of EMG envelopes as:

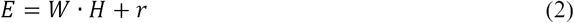

where E is a *m* × *t* envelope matrix (m – number of muscles, t – number of samples used); W is a *m* × *s* synergy vectors matrix (s – number of modules) containing the relative weights of each muscle within each synergy; H is a *s* × *t* activation pattern (AP) matrix; r is a root mean square residual error that describes difference between reconstructed envelope matrix and original one. The NNMF algorithm was applied to epochs consisting of consecutive bouts of 10 cycles concatenated together. The quality of reconstruction was measured using the variance accounted for (VAF):

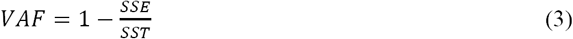

where SSE is sum of squared errors and SST is total sum of squares. (Torres-Oviedo et al. 2006). We removed the mean values from the EMG envelopes when calculating SST. Muscle synergies were extracted unilaterally, thus analysing separately the muscles of the left and right legs.

Previous studies in literature have found that cycling is well described by using either 4 or 5 unilateral modules (Barroso et al. 2014; De Marchis et al. 2013; De Marchis et al. 2015). We performed synergy analysis on the muscles of each leg by extracting 4 and 5 unilateral modules and selected the most appropriate reconstruction for further analysis as the one yielding an average VAF across subjects > 90% and > 80% for each muscle.

### Analysis of the changes in muscle synergy activation patterns and in the EMG activity

As changes in the accelerometer were characterized by a shift in the peak of the acceleration of the crank arm, we assessed for temporal shifts of the APs during the course of the two experiments. This analysis was based on the calculation of the lag of the maximum of the cross-correlation function calculated between the average AP of each synergy extracted during the whole BL and the average APs extracted from each epoch. We also compared, for each synergy extracted from each leg, the average lag calculated at late BL, early AD, late AD and early PA. In this analysis changes were evaluated, across the different sections of both experiments, using Friedman’s test (with Bonferroni’s correction with number of comparisons equal to the number of synergies and thus of tests performed on each leg) and a post-hoc analysis based on Dunn-Sidak’s criterion (non-normality of the data was assessed using the Kolmogorov-Smirnov test).

Changes in the acceleration patterns due to the asymmetries could also be caused by modifications in the activation level of the different muscles during a cycle. To capture this, we assessed changes in the amount of the muscular activity of all 16 muscles in the two experiments. This analysis was based on the calculation of the integrated EMG (iEMG) from the average envelope of each muscle in each epoch. Changes in iEMG were expressed as percentage changes of the iEMG calculated in each epoch with respect to its average value during the whole BL phase. Consistently with the lag analysis, we investigated the percentage changes in iEMG through all the epochs and on the average values calculated at late BL, early AD, late AD and early PA. Also in this case changes were evaluated, across the different sections of both experiments, using Friedman’s test (with Bonferroni’s correction with number of comparisons equal to the number of muscles on each leg) and a post-hoc analysis based on Dunn-Sidak’s criterion (non-normality of the data was assessed using the Kolmogorov-Smirnov test).

## Results

All accelerometer and EMG data were checked for quality before analysis. Subjects with poor EMG or accelerometer data (that is, data that presented either discontinuities due to wireless disconnections or non-physiological artefacts such as non-movement related spikes) were excluded from the final analyses. After this assessment, three subjects were dropped in the analysis of the accelerometers for both experiments, while one subject was excluded from the muscle synergy analysis of the Angle experiment and two subjects were excluded from the analysis of the Length experiment. None of the subjects reported exhaustion or fatigue at the end of each phase of the experiments.

### Accelerometers

The results of the analysis of the accelerometer data for the Angle and Length experiments respectively are presented in **Fig. 2** and **Fig. 3**. In both experiments we observed shifting patterns in the profile of the sagittal acceleration of the left crank arm during the different phases of the experiments (**Figures 2A** and **3A**). Changes in magnitude in the acceleration were observed in the first two to five cycles of all three phases (as the subjects reached the required pace/power output) of the experiment and were consistent across all phases and experiments (as exemplified by the plot showing the 1^st^ cycle of AD in **Figures 2A** and **3A**). These behaviours were likely related to the longer cycle duration as subjects accelerated during the first cycles of all phases of both experiments (**Figures 2D** and **3D**).

**Fig 2.**
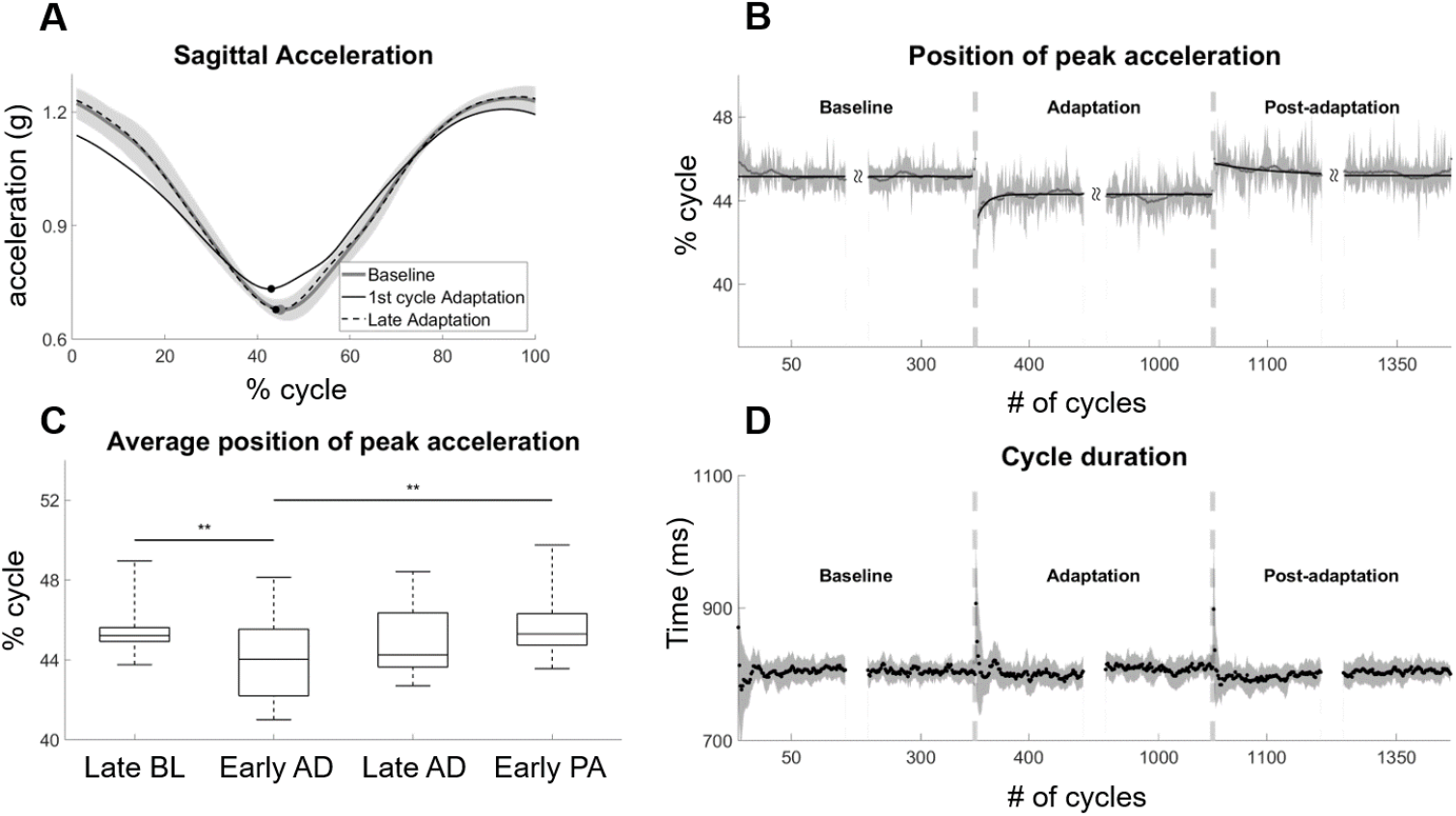
Changes in the profile of the sagittal acceleration – Angle experiment. Adaptation in the acceleration was characterized by tracking the changes in the position of the peak sagittal acceleration. *A* presents the profile of the sagittal acceleration at different points during the experiment. The shaded area represents the average (grey solid line) ± standard deviation (across subjects) of the average sagittal acceleration recorded during BL. The black solid line represents the sagittal acceleration in the first cycle of adaptation while the black dashed line represents the average sagittal acceleration at late adaptation (last 50 cycles). Bold markers represent the position of the minima for the three curves. In *B* the shaded area represents the median (grey solid line) ± standard error values of the position of peak sagittal acceleration through all the cycles of the experiments. The black bold lines represent linear/exponential fitting (in the form presented in equation 1) of the median data. In *C* each boxplot represents the median, 25th and 75th percentiles, minimum and maximal values of the position of the peak sagittal acceleration across the subjects on the data averaged across different phases of the two experiments. Late BL and Late AD refer to the last 50 cycles of BL and AD while Early AD and Early PA refer to the first 50 cycles of AD and PA. ** indicates statistical significance with p<0.01 according to Dunn-Sidak’s method. In *D* is presented the duration (in ms) of each individual cycle. Vertical dashed lines represent the beginning of the AD and PA phases.

**Fig 3.**
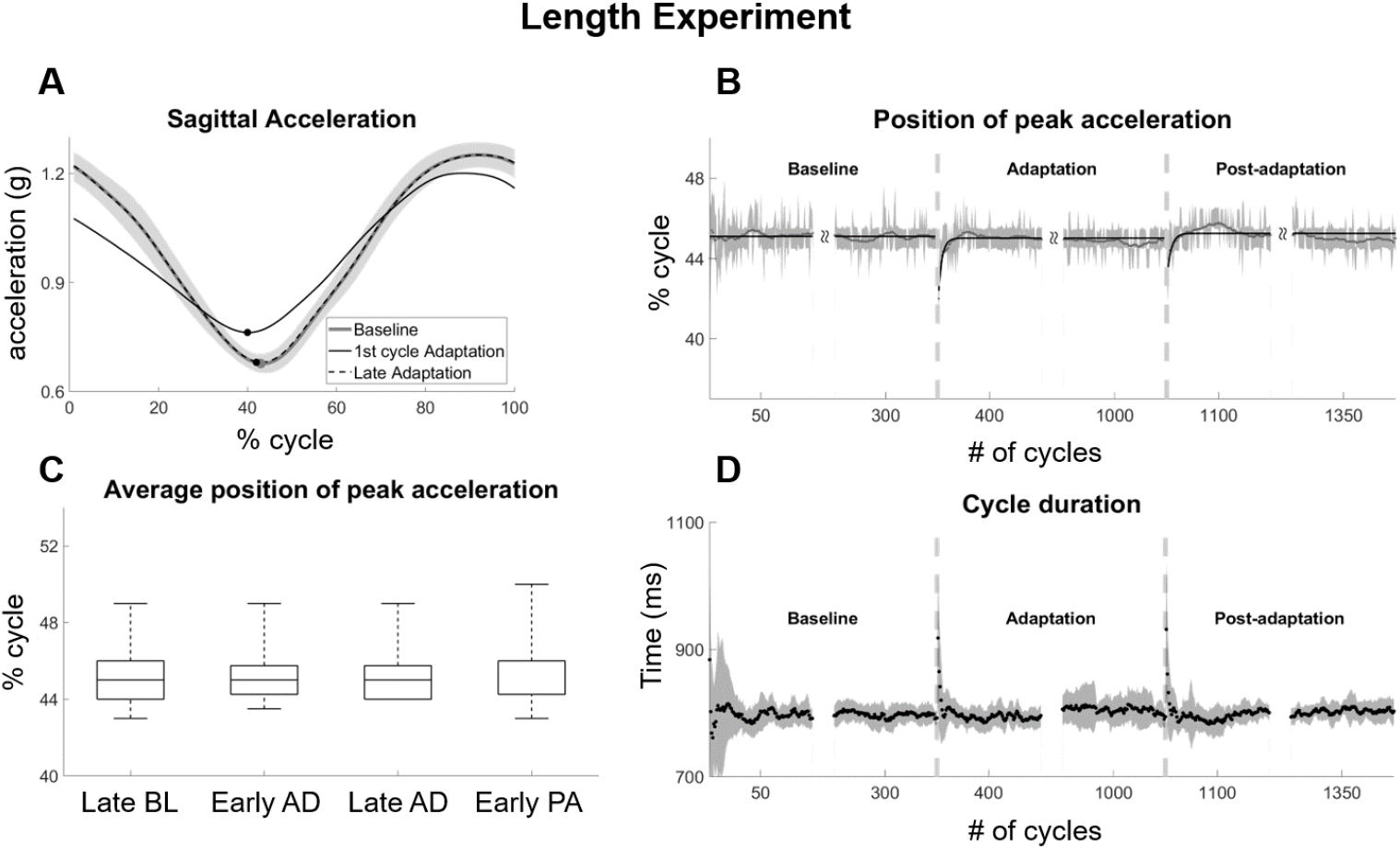
Changes in the profile of the sagittal acceleration – Length experiment. The organization of this figure is identical to that of Fig. 2.

In the Angle experiment we observed a shifting pattern in the profile of the accelerometer data. This behavior was well captured by tracking the position of the minimum of the sagittal acceleration pattern for each cycle and is consistent with an exponential adaptation and washout. The position of the peak acceleration was observed to be stable around a constant value (45.2 ± 1.6 % of the cycle) during the BL phase of the experiment (**Figure 2B**) and to shift earlier in the cycle at the beginning of the AD phase (41.7 ± 2.8 % in the first cycle of AD). The peak of acceleration progressively shifted back in the cycle until it reached a plateau at about 44.5 % of the cycle towards the end of the AD phase. This behavior can be modelled by an exponential function in the form presented in equation (1) and with a time constant (intended as the number of cycles that are necessary to reach the plateau from the beginning of adaptation) equal to 25 cycles (see **Table 1**). At the beginning of the PA phase we observed an after-effect opposite in direction with respect to the original peak shift observed at the beginning of AD. This translates in a shift forward in the peak of the acceleration up to about 46.0% of the cycle. We observed a slow exponential behavior (time constant equal to 166 cycles) related to the washout of the adapted behavior. A non-parametric Friedman test was conducted to assess for statistically significant changes in the peak of acceleration across the different phases of the experiment. The test rendered a Chi-squared value of 15.24 which was significant (p<0.01). Post-hoc analysis, based on Dunn Sidak’s test, showed that the position of the peak in early AD was statistically different than its corresponding values at late BL (p < 0.01) and early PA (p < 0.01).

**Table 1.** Exponential fitting parameters for position of the peak sagittal acceleration for the AD and PA phases of the Angle and Length experiments. All values are presented together with ± 95% confidence intervals.

| | | $\alpha$ | $\beta$ | $\gamma$ | Time Constant |
| --- | --- | --- | --- | --- | --- |
| **ANGLE** | **AD** | -1.43 (-1.58, -1.27) | 0.120 (0.102, 0.138) | 44.31 (44.3, 44.32) | 25.00 (21.74, 29.41) |
|  | **PA** | 0.59 (0.55, 0.64) | 0.018 (0.015, 0.021) | 45.20 (45.17, 45.22) | 166.67 (142.86, 200.00) |
| **LENGTH** | **AD** | -3.49 (-3.71, -3.27) | 0.285 (0.263, 0.307) | 45.01 (45.01, 45.02) | 10.52 (9.77, 11.41) |
|  | **PA** | -2.51 (-2.91, -2.11) | 0.259 (0.206, 0.312) | 45.25 (45.22, 45.27) | 11.58 (9.62, 14.56) |

In the Length experiment we again observed an exponential shift in the peak of the sagittal acceleration (**Figure 3B**) during the adaptation phase. As for the Angle experiment, BL was characterized by consistent values of peak acceleration (45.2 ± 1.6 % of the cycle). At the beginning of the AD phase we observed an initial anticipation in the peak value of acceleration (42.4 ± 7.0 % in the first cycle of AD) that was exponentially shifted back close to the BL value by the end of AD. The time constant for this exponential behavior was estimated to be equal to 10 cycles (**Table 1**). At the beginning of PA, instead of an after-effect opposite to the original error at the beginning of AD as expected in a motor adaptation model and as observed in the Angle experiment, we observed once again a backward shift in the peak of the acceleration, in a behavior almost identical to the one observed during the AD phase. This exponential behavior was characterized by a time constant of about 11 cycles, similar to the one observed in the AD phase (**Table 1**). Statistical analysis on the first/last 50 cycles of the different phases did not show statistically significant differences in the average position of the peak acceleration (Friedman’s test, Chi-squared value = 1.79, p = 0.62), as the initial changes in peak position at the beginning of AD and PA were not captured in this analysis due to the shortness of the relative exponential behaviors (10 cycles, against a window of analysis of 50 cycles) and to the fact that the initial shift was perfectly compensated for. It is possible that by shortening the window of analysis to 10 cycles we would have observed statistically significant changes in position between early and late AD. Nevertheless, we decided to keep the window of observation consistent across the different analyses to make the results as much comparable as possible between the different measures (accelerometer, EMG and synergies).

#### Muscle Synergy Analysis

. In our analysis we compared the quality of EMG reconstruction after extracting either 4 or 5 synergies (**Figure 4A**), that are model orders commonly used in synergy analysis during cycling (Barroso et al. 2014; De Marchis et al. 2013). We found that 4 synergies yielded VAF values of 84.0% for the Angle experiment and 84.9% for the Length experiment, while 5 synergies values of VAF equal to 91.3% and 91.9% respectively for the two experiments. Based on these results we decided to perform all further analysis on the 5 synergies reconstruction. The synergy sets we extracted (**Figure 4B**) are in line with those found in similar works in literature (Barroso et al. 2014; De Marchis et al. 2013; De Marchis et al. 2015) and are constituted by the following modules: Synergy 1 is constituted by the quadriceps muscles (RF and VL) and reflects hip flexion and knee extension in the first quadrant of the cycle (taking as starting point the highest position of the pedal during the cycle); Synergy 2 is constituted by the muscles of the calf (Sol and GM) and controls ankle dorsiflexion in the second quadrant of the cycle; Synergy 3 is mostly constituted by the activity of the BF and GM and is also active in the second quadrant, controlling ankle dorsiflexion and knee extension; Synergy 4 is constituted by the TFL and controls hip flexion in the third quadrant; Synergy 5 is constituted by the TA and is mostly active in the fourth quadrant and is relative to ankle plantarflexion.

**Fig 4.**
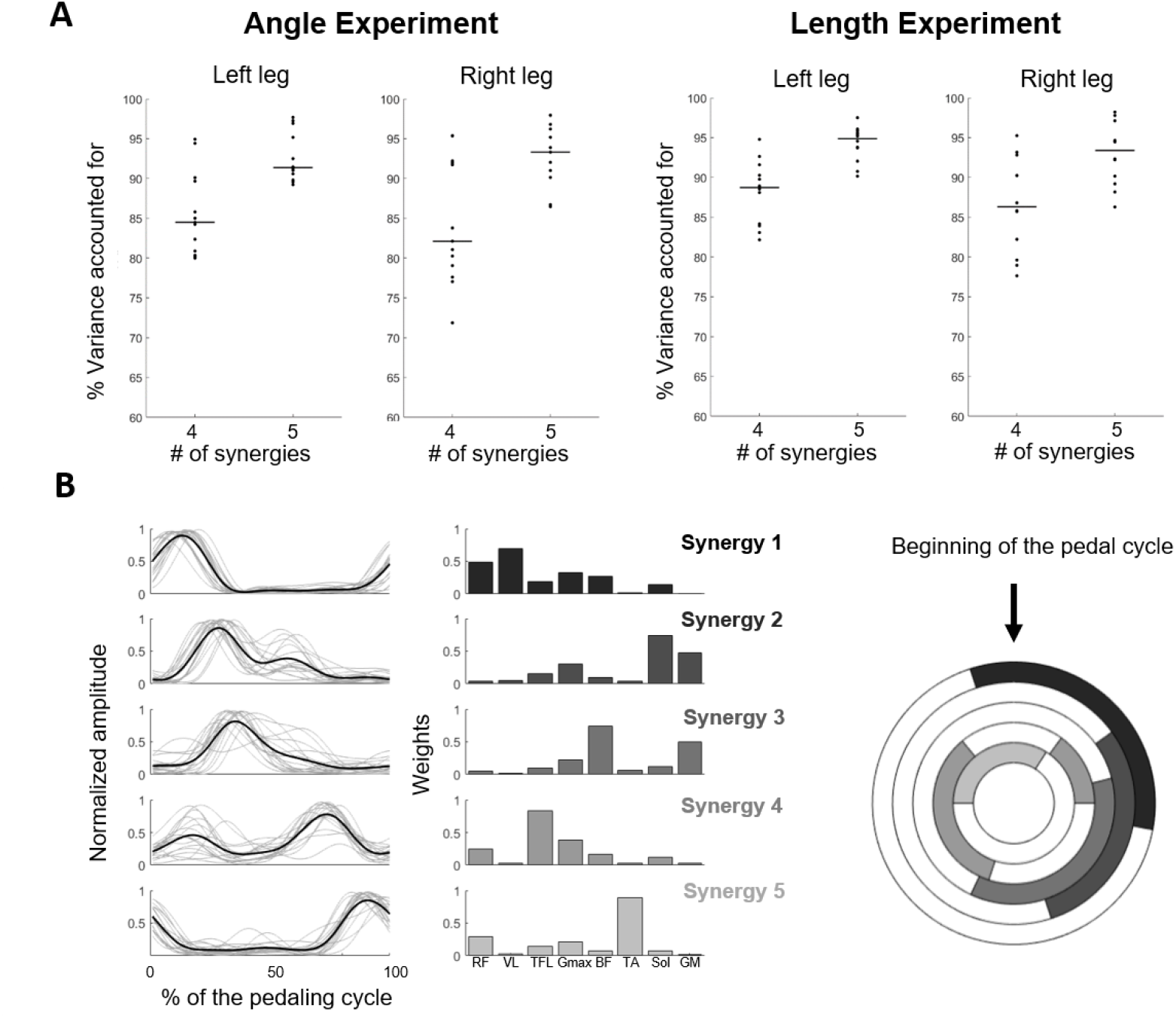
Muscle Synergies Extraction. In *A* are shown the values of VAF obtained by extracting either 4 or 5 synergies from the data recorded during the baseline phase of each experiment of each subject for both the left and right leg. Individual dots represent the average VAF across the muscles of each leg, while horizontal lines represent average values across subjects. In *B* are shown the activation patterns (bold lines represent the average across subjects, light lines represent individual subjects) and the average weights recorded across subjects. On the right are presented the activation patterns as mapped in a circle representing the position of the foot during a cycle (with the top position representing the beginning of the pedal cycle, marked as the bottom of the cycle on the left foot). For this representation, a hard threshold of 0.2 for the average APs (normalized to their maximum value) was used to determine the onset and offset of the activation in the cycle.

#### Changes in Synergy Modules and Activation Patterns

We qualitatively analysed the changes in synergies modules and APs through the different phases of the two experiments to capture specific neuromuscular behaviors consistent with the biomechanical changes we observed in the analysis of the accelerometers. The composition of the 5 motor modules remained consistent in both experiments across the different phases (**Figure 5**). In the Angle experiment we observed small fluctuations in the weightings of some muscles on the synergies of both the left (e.g. Gmax and Sol in Synergy 2, GM in Synergy 3) and the right sides (e.g. Sol and GM in Synergy 2, BF in Synergy 3). Similarly, in the Length experiments we observed small bilateral changes (e.g. Sol in Synergy 2 and TA on Synergy 5 on the left side, GM in Synergy 2 and 3 and TA in Synergy 5 on the Right side).

**Fig 5.**
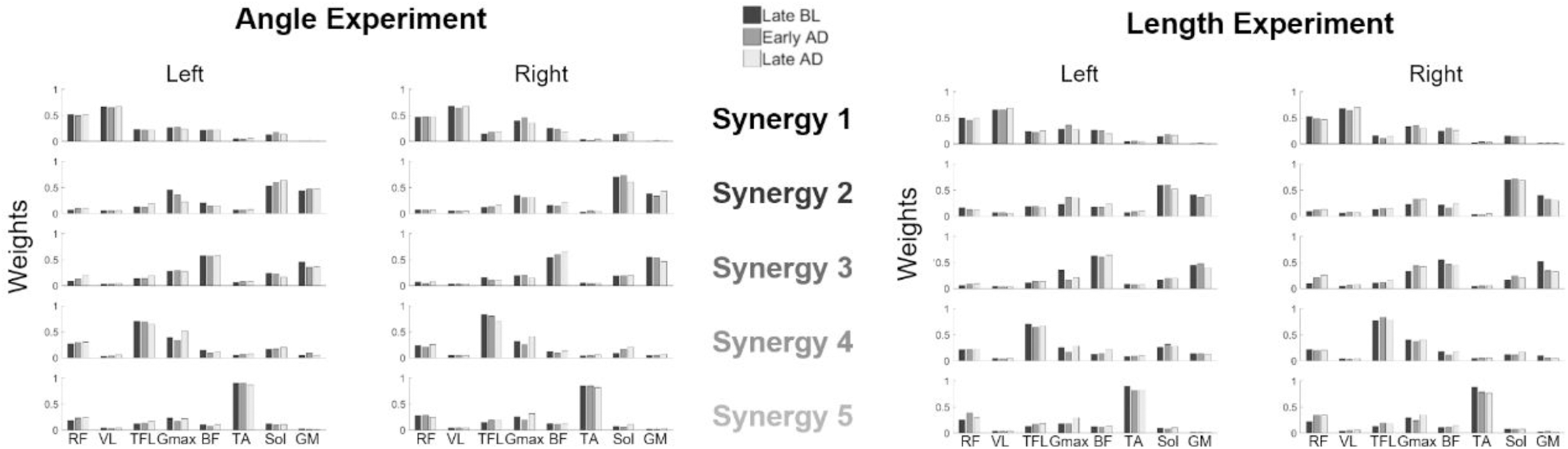
Synergy modules across the different phases of each experiment. Each bar represents the average (across subjects) value of the contribution of a muscle to each synergy (color coded as in Fig. 4) estimated across the different phases of the two experiments. Black represents Late BL, grey Early AD and light grey Late AD.

In both experiments we observed qualitative changes in the APs of the synergies modules that were mostly characterized by small changes in amplitude and peak timing in the cycling pattern (**Figure 6A** and **6B**). In the Angle experiment, in a visual analysis of **Fig. 6A,** we noted progressive changes in APs amplitude in synergies 2, 3 and 4 on the left side and in all synergies on the right side. Moreover, we observed progressive shifts in the activation timing of the APs of Synergies 2 and 3 for the right side. These changes reflect a progressive anticipation of the activity of the calf muscles and of the BF during the cycle that possibly translates into an anticipation of ankle plantar flexion and knee flexion during the cycle, necessary to compensate the anticipation in peak acceleration. In the Length experiment we again observed progressive changes in synergy amplitude on most of the synergies on both sides, but we did not observe clear shifts in the position of the APs. In order to capture quantitatively the shifting of the APs we calculated the lag of the cross-correlation functions between the average APs during BL and those calculated in all the epochs of the experiment (with particular focus on late baseline, early and late adaptation and early post-adaptation). In the Angle experiment (**Figure 7**) we did not observe shifts in the position of the APs for the synergies of the left leg, with median values of lags around 0 through all phases of the experiment for all the 5 synergies. On the right leg, however, we observed clear progressive shifts in AP position for synergies 2 and 3, as already qualitatively observed from **Fig. 6A**. For these synergies we observed a gradual anticipation of the APs during the AD phase that quickly stabilizes around a plateau value and is quickly washed out at the beginning of the PA phase.

**Fig 6.**
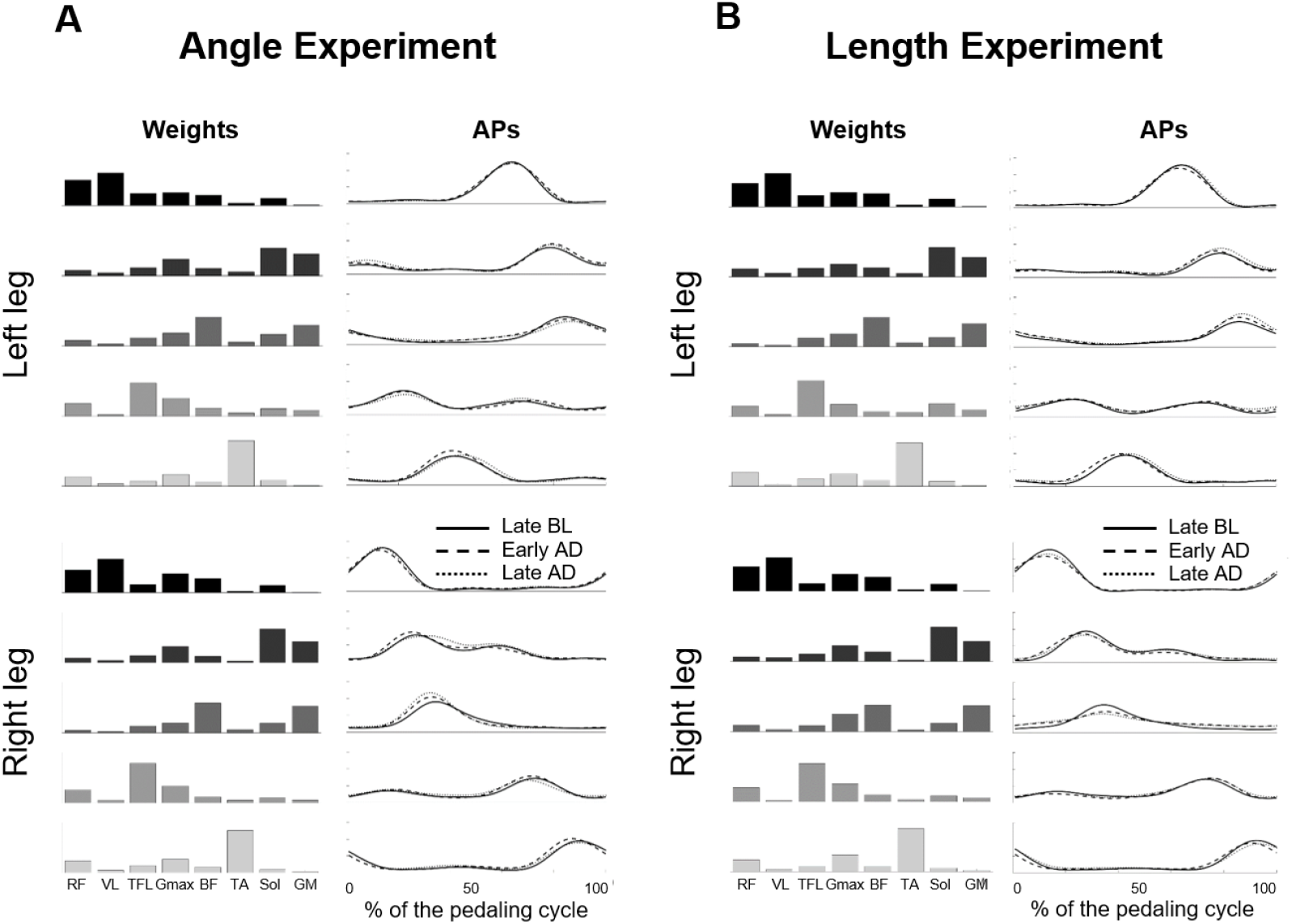
Weights and APs of the muscle synergies across the different phases of the experiments. Panel *A* presents the results of the Angle experiment, panel *B* of the Length experiment. For each panel, top plots represent the left leg, while bottom plots represent the right leg. Synergies modules are the average modules at baseline and are showed for reference. For each AP plot, the bold line represents Late BL, the dashed line represents Early AD and the dotted line represents Late AD.

**Fig 7.**
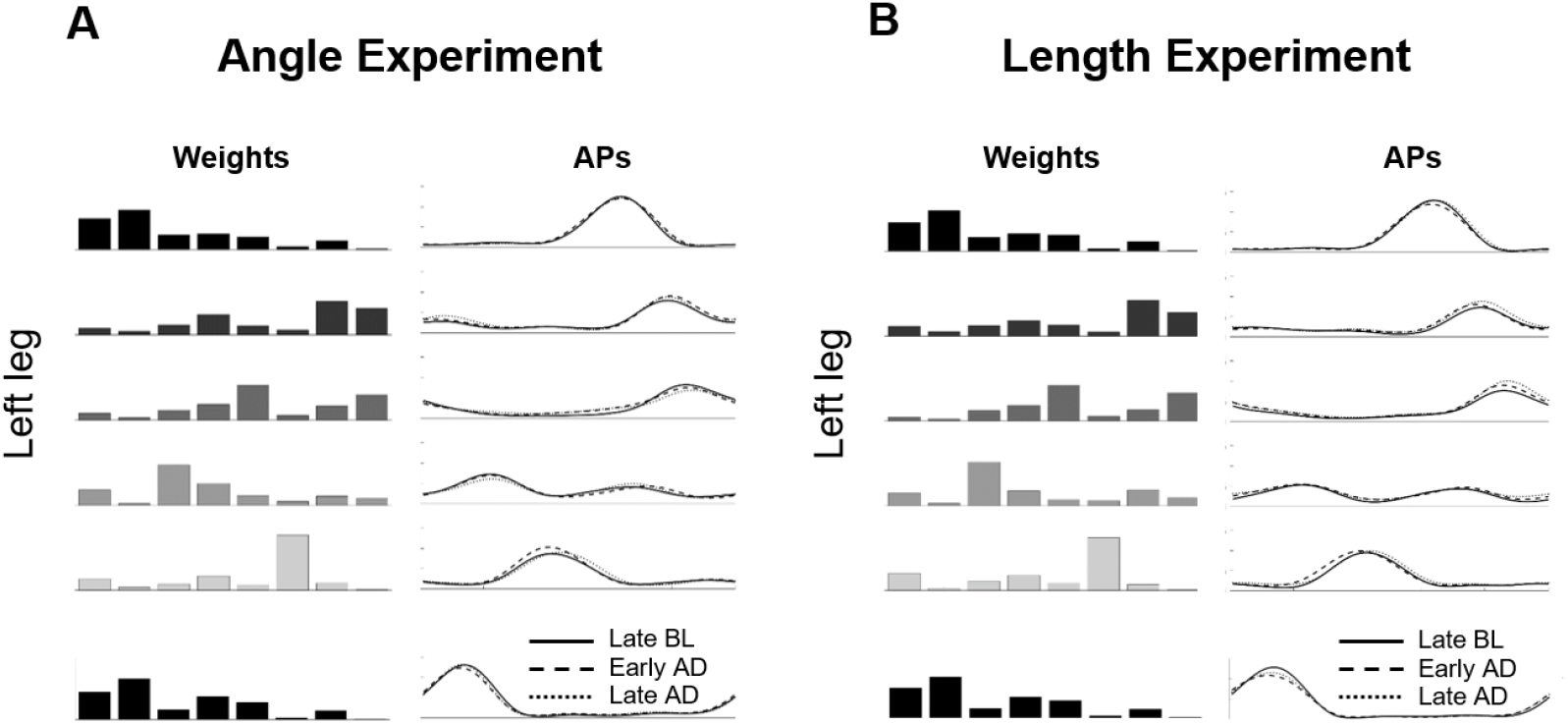
Temporal shifts in the APs of the muscle synergies – Angle Experiment. The column on the left for each leg presents the lag between the average baseline APs and the APs of each epoch, through all the experiment. Each point represents the median lag across subjects in one epoch, while the shaded area represents the standard error. The two vertical lines represent the beginning and end of the AD phase. The right column represents the average and standard deviation of the lag calculated during Late BL, Early AD, Late AD and Early PA. * indicates statistically significant differences with p<0.05 (based on Dunn-Sidak’s method). Each row represents a muscle synergy, color-coded as in Fig. 4.

Statistical analysis confirmed these observations. On the left side, the results of the Friedman’s tests performed on the average lag calculated in late BL, early and late AD and early PA for all synergies did not result in the observation of statistically significant changes (Chi-squared 2.26, p = 0.52 for Synergy 1, Chi-squared 2.04, p = 0.56 for Synergy 2, Chi-squared 3.64, p = 0.30 for Synergy 3, Chi-squared 4.05, p = 0.26 for Synergy 4, Chi-squared 1.79, p = 0.62 for Synergy 5). On the right side, we observed statistically significant differences in lag in Synergy 2 and Synergy 3 (Chi-squared 9.7, p = 0.02 for Synergy 2, Chi-squared 10.71, p = 0.01 for Synergy 2). Post-hoc analysis revealed statistically significant changes between late AD and early PA in Synergy 2 (p = 0.018) and between late BL and both early and late AD in Synergy 3 (p =0.027 and p = 0.043 respectively). The other synergies on the right side did not present statistically significant changes (Chi-squared 6.97, p = 0.07 for Synergy 1, Chi-squared 2.25, p = 0.52 for Synergy 4, Chi-squared 3.39, p = 0.33 for Synergy 3). In the Length experiment (**Figure 8**) we did not observe shifts in the AP of all the synergies on both the left (Chi-squared 3.15, p = 0.37 for Synergy 1, Chi-squared 1.24, p = 0.74 for Synergy 2, Chi-squared 4.53, p = 0.21 for Synergy 3, Chi-squared 1.11, p = 0.77 for Synergy 4, Chi-squared 1.79, p = 0.62 for Synergy 5) and right sides (Chi-squared 3.24, p = 0.36 for Synergy 1, Chi-squared 2.69, p = 0.44 for Synergy 2, Chi-squared 0.43, p = 0.93 for Synergy 3, Chi-squared 0.84, p = 0.84 for Synergy 4, Chi-squared 5.7, p = 0.12 for Synergy 5).

**Fig 8.**
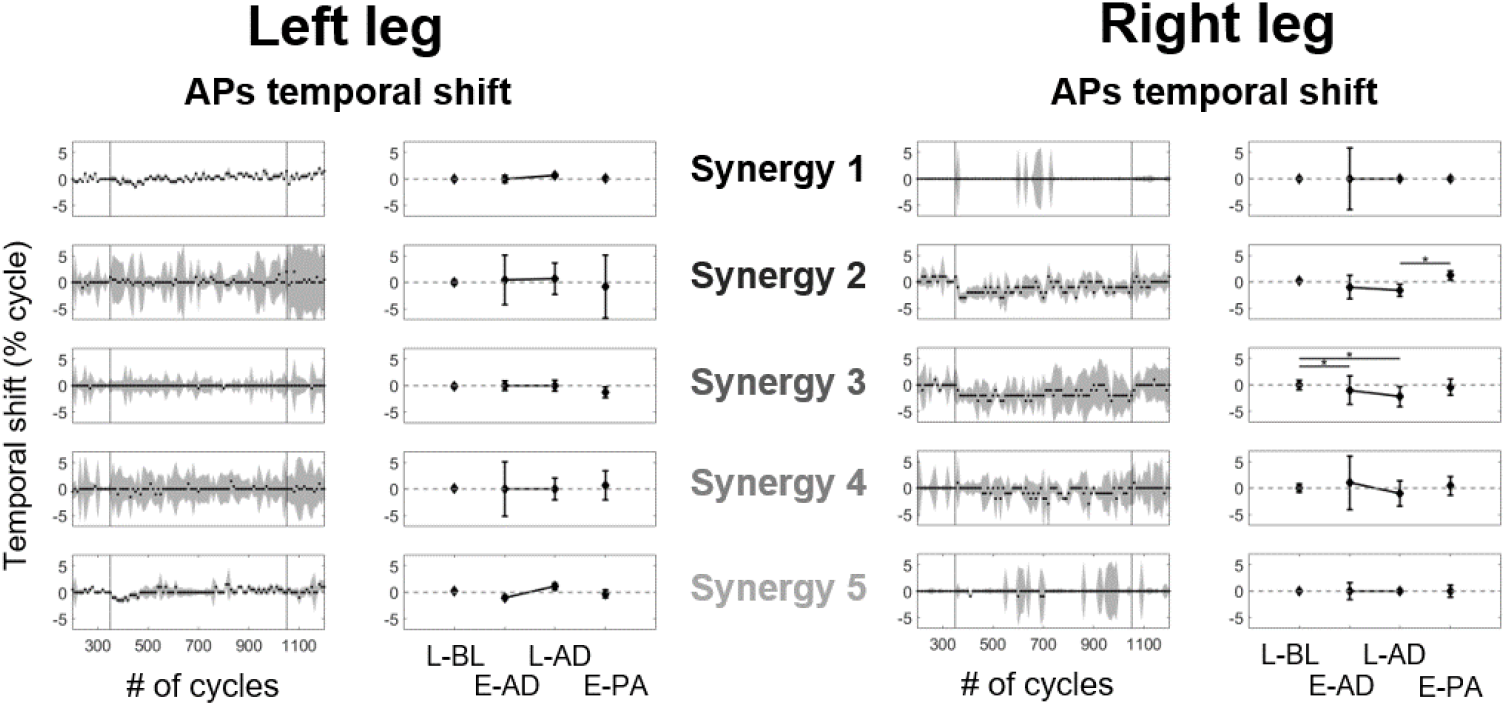
Temporal shifts in the APs of the muscle synergies – Length Experiment. The organization of this figure is identical to that of Fig. 7.

#### Changes in EMG activity

As a final analysis we investigated if the introduction of the two asymmetries induced changes in the amount of EMG activity in all of the 16 muscles under analysis. This analysis was based on the calculation of the percentage changes (with respect to its average value at BL) of the iEMG calculated from each epoch. In the Angle experiment the introduction of the asymmetry did not correspond with clear increases or decreases in muscular activity (**Figure 9**). We observed a trend of increase in EMG activity in the left Sol and in the right GMax, BF and Sol (these muscles also presented increased variability). However, none of these muscles presented statistically significant changes in iEMG across the different phases of the exercise (Chi-squared and p-values omitted for brevity). In the Length experiment (**Figure 10**), once again, we did not observe clear changes in iEMG across the different muscles of the left leg, and all comparisons were shown to be not significant (Chi-squared and p-values omitted for brevity). In the right leg, however, we observed a clear decrease in iEMG activity for the GM muscle that corresponded exactly with the AD phase. These changes were shown to be statistically significant (Chi-squared = 22.24, p < 0.01). Post-hoc analysis revealed statistically significant differences between iEMG calculated at late BL and both early and late AD (p < 0.01 in both cases) and between early PA and both early and late AD (p = 0.014 and p < 0.01) respectively. None of the other muscles on the right leg showed statistically significant differences in iEMG across the different phases of the experiment (Chi-squared and p-values omitted for brevity).

**Fig 9.**
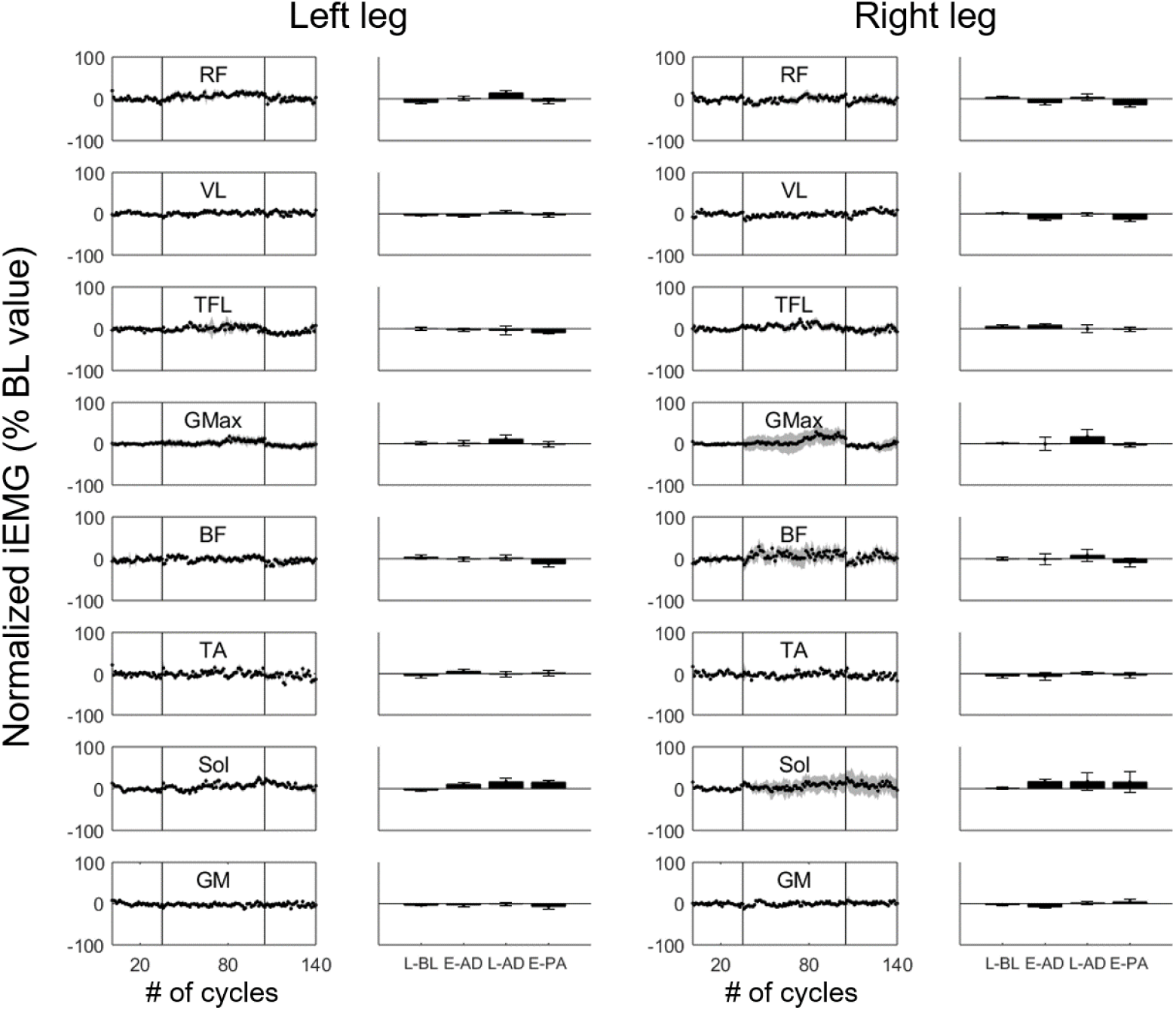
Changes in iEMG – Angle Experiment. The column on the left for each leg presents the value of iEMG for each epoch, through all the experiment. Each point represents the median lag across subjects in one epoch, while the shaded area represents the standard error. The two vertical lines represent the beginning and end of the AD phase. The right column represents the average and standard deviation of the iEMG calculated during Late BL, Early AD, Late AD and Early PA.

**Fig 10.**
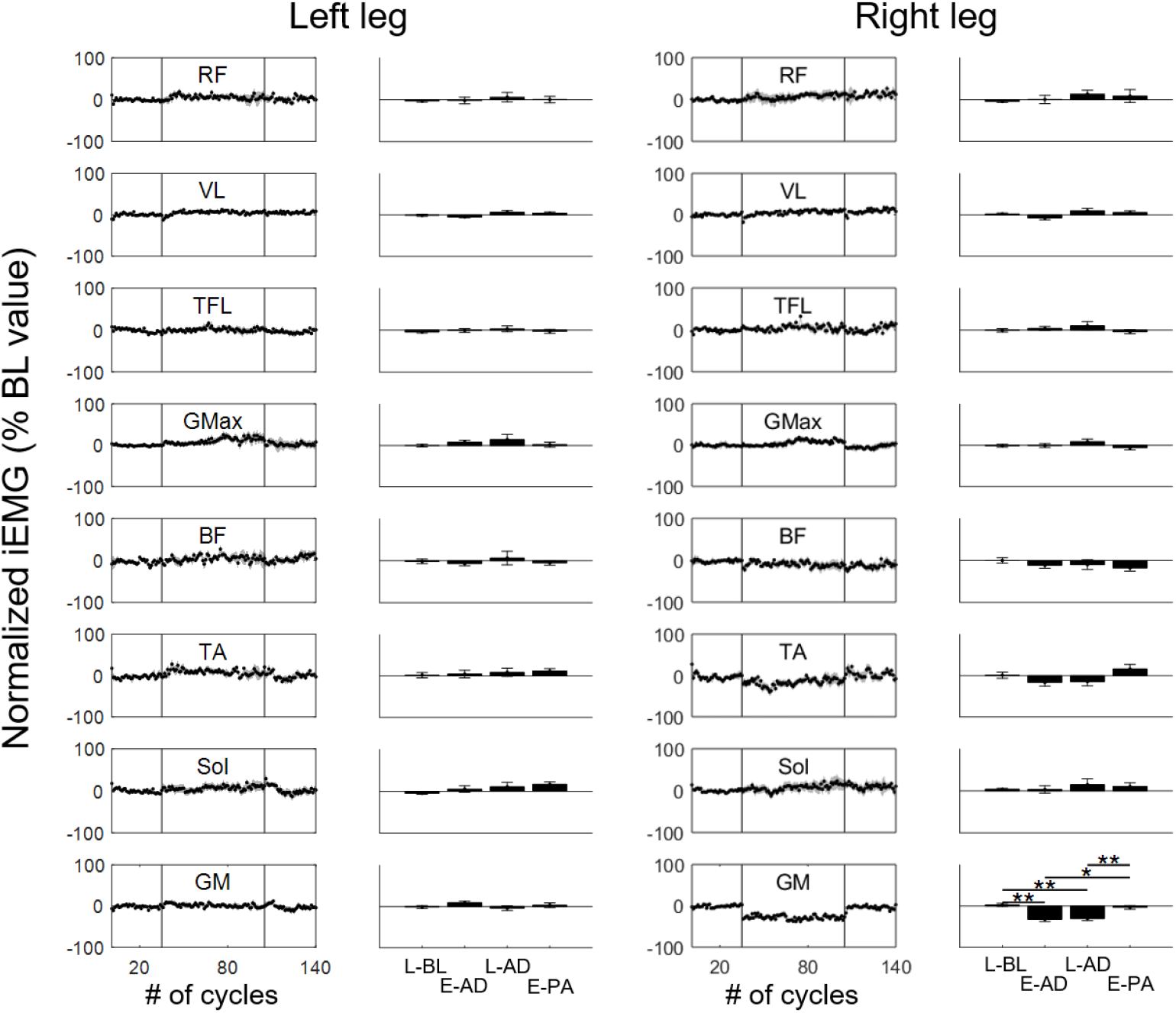
Changes in iEMG – Length Experiment. The organization of this figure is identical to that of Fig. 9. * indicates statistically significant differences with p<0.05 (based on Dunn-Sidak’s method), while ** indicates statistically significant differences with p<0.01.

### Discussion

In the two experiments herein presented, we investigated if small asymmetries in the angle and length of the crank arms of the pedals during cycling translate into motor adaptations. We found that altering the angle between the two pedals leads to a progressive shift in the crank acceleration that is consistent with a motor adaptation, intended as a progressive update of the motor plan, and presents an opposite after-effect once the asymmetry is removed. This adaptation appears to be mainly obtained by modulating the timing of activation of some of the synergies on the “perturbed” side (herein indicating the right side, where the asymmetries were introduced, although we acknowledge that, at least for the angle experiment, it is not possible to strictly define a perturbed side). For the length asymmetry, we observed a behavior characterized by a fast exponential response that is equivalently (in magnitude, timing and side) observed in both adaptation and post-adaptation. At the neuromuscular level, for the Length experiment, we did not observe changes in the timing of the muscle synergies. However, we observed a sudden step-like decrease in the overall activity of the right GM muscle as the asymmetry is introduced. The activity of the muscle returned to normal once the asymmetry was removed. The lack of an opposite after-effect during post-adaptation in the accelerometer data and the lack of an adaptive behaviour (intended as an exponential change) in the neuromuscular data suggest the compensations observed in the Length experiment are not consistent with a predictive feedforward motor adaptation.

### Angle experiment, insights into possible adaptation mechanisms

In the Angle experiment we observed an adaptation-like behavior that is reflected, at the biomechanical level, by a shift in the peak of the sagittal acceleration of the crank arm. This adaptation converges in about 25 cycles, and is characterized, at the neuromuscular level, by a similarly timed exponential shift in the activation of the 2 synergies on the perturbed side, whose activation patterns peak at the dead bottom of the cycle. The adaptation that we observe for the Angle experiment produces a small after-effect at the biomechanical level after the asymmetry is removed (Figure 2), while no step-like changes are observed at the neuromuscular level at the beginning of PA. The lack of step-like changes in the muscular activity when the asymmetry is removed confirms that the biomechanical aftereffect is caused by an adaptation of the motor plan. In fact, when the asymmetry is removed the motor plan is optimized for rejecting the perturbation but not for “unperturbed” cycling. This discrepancy causes the biomechanical aftereffect and triggers the washout (that is, in fact, a re-optimization of the motor plan to the unperturbed condition) that we observed both at the biomechanical (Figure 2) and neuromuscular (Figure 7, synergies 2 and 3) levels. It is worth noticing that the exponential expression of motor adaptations is caused by the progressive decrease in biomechanical error driven by the cycle-by-cycle update of the internal models driving the task (Kawato 1999; Scheidt et al. 2001; Shadmehr and Mussa-Ivaldi 1994). In the Angle experiment the asymmetry introduces a fixed static alteration in the half-cycle relationship between the two legs (Ivanenko et al. 2004), but since the position of the crank arm cannot be altered, the kinematic error cannot be compensated over the course of the exposure to the experiment. Thus, our results point towards the presence of implicit kinetic or temporal error measures that are minimized during the AD phase. These error measures are likely driven by force and position proprioceptors (Jensen et al. 1998), that map the relative timing between the hypothetical CPGs driving the two legs similarly to what has been observed in cats (Frigon et al. 2013). This result is not unexpected as adaptation specific to temporal parameters of repetitive leg movements was also observed during split-belt treadmill experiments in humans (Malone et al. 2012). The adaptation that we observed in the neuromuscular activity of the subjects during the Angle experiment appears to be circumscribed to the “perturbed” leg. In fact, we did not observe bilaterally-linked temporal adjustments in the muscle synergies as previously observed in the split-belt treadmill experiment (MacLellan et al. 2014). Our results seem more in line with side-specific adjustments in the timing of the CPGs (Choi and Bastian 2007) that is, however, driven by a bias toward a fixed synchronization between the two legs (Reisman et al. 2005).

### Length experiment, potential influence of spinal reflexes

In the Length experiment we observed, at the biomechanical level, an initial anticipation in the peak of the sagittal acceleration of the crank arm that is quickly (time constant equal to 10 cycles) shifted back to its baseline value and does not present an opposite after-effect but an almost identical behavior also during PA. At the neuromuscular level, as expected, we did not observe noticeable temporal adjustments in the synergies APs (**Figure 8**), as the timing between the two legs had been left unaltered. We instead observed a localized response that was characterized by a step-like decrease in activity in the gastrocnemius medialis muscle during the AD phase. This decrease was equally discarded in a step-like manner once the asymmetry was removed (**Figure 10**). The behavior we observe in the Length experiment is more consistent with a feedback response rather than a feedforward motor adaptation. Locomotor adaptations have been shown to be constituted by both feedback and feedforward components (Lam et al. 2006; Reisman et al. 2005). The former are fast adjustments that do not yield an aftereffect once the perturbation is removed (as in the case of the GM in the Length experiment), while the latter are slower adjustments that are progressively learned, consistently with the update of a motor plan, and produce an after effect in the opposite direction (as we observed in the acceleration patterns for the Angle experiment). In this interpretation in the Length experiment we observe a purely feedback response to the asymmetry that is localized in the GM muscle and that is quickly discarded as the asymmetry is removed. This response is likely caused by the difference in task demands caused by the decreased pedal length. Nevertheless, how the neuromuscular changes we observed in the GM muscles correlate with the changes in the acceleration patterns remains elusive.

### Synergistic representation of feedback and feedforward responses

Although the muscle synergies modules identified in our analysis maintain a noticeable consistency between the two experiments and across the different phases of both experiments (**Figure 5**), we observed different overall behaviors in the synergistic representation of the feedforward and feedback mechanisms observed in the two experiments. In the Angle experiment the neuromuscular changes due to the asymmetry are exclusively observed in the activation patterns of the synergies while the shapes of the modules remain consistent through the different phases of the experiments. These results are in accordance with the majority of previous literature that has shown that synergies modules are robust across different tasks and task demands. During lower limb movements, synergy modules have been shown to be robust through different walking conditions (Chia Bejarano et al. 2017; Courtine et al. 2006; Oliveira et al. 2016), balance perturbations (Oliveira et al. 2013; Torres-Oviedo et al. 2006), cycling conditions (De Marchis et al. 2013; Hug et al. 2011), fatigue (Castronovo et al. 2018; Turpin et al. 2011) and to be shared across different tasks (Barroso et al. 2014; Chvatal and Ting 2013). Similar results have also been observed during motor adaptations. Studies on visuomotor rotations in the upper limbs have demonstrated that APs are rotated while modules remain consistent when adapting to different visual perturbations (Berger et al. 2013; De Marchis et al. 2018; Gentner et al. 2013) and similar results have been also observed during split-belt treadmill experiments (MacLellan et al. 2014). Our results seem to suggest a rotation effect in the APs of the muscle synergies during the Angle experiment similar to the one observed during visuomotor rotations (De Marchis et al. 2018; Gentner et al. 2013). It was indeed expected that a potential adaptation to the temporal asymmetry would have been explained through a modification of the timing of the APs.

On the other hand, in the Length experiment, it was expected that the asymmetry would alter the relative contribution of the different muscles to the task. In fact, Ranganathan et al. (Ranganathan et al. 2016) showed, in a conceptually similar task performed while walking in the Lokomat device, that adjustments to different kinematical demands can be obtained by changing the modular structure of the synergies. Indeed, we observed a decrease in activity only in the GM muscle during AD (**Figure 10**), that is reflected in the small fluctuations in the weight of this muscle in Synergies 2 and 3 (**Figure 5**). Interestingly, these changes are evident and significant only for GM, although also the TA, the antagonist of GM, shows a qualitative decrease in activity. However, the decrease in activity that we observed in the GM muscle is not paired with a similar decrease in its main synergistic companions (Sol in Synergy 2 and BF in Synergy 3), indicating that such behaviour cannot be modelled as the decrease in activation of one or more synergies. The results of the Length experiment could suggest then that the contribution of the different muscles to the synergies can be slightly modulated in response to altered task demands. This could be caused by a modulation in the second layer of the CPGs, that supposedly controls muscular pattern formation (Zhong et al. 2012). A perhaps more plausible explanation of our results could be that the shorter pedal length induces a change in the length of the GM muscle during its force-production phase and that the decrease in activity of the muscle is due to an adjustment of its activation in response to the different length-tension relationship.

### Conclusions

The results herein presented show that feedforward motor adaptations in the lower limbs are present in response to small temporal asymmetries during cycling, while spatial asymmetries elicit feedback compensations. The adaptations we observe are present when balance is not directly compromised and when the kinematic error cannot be compensated for. Our results suggest that the neural circuits controlling the timing of the legs during repetitive movements adapt to maintain specific symmetry features in the unilateral muscular activation patterns. Moreover, we show that synergies modules can be modified in response to altered task demands.

### Limitations

There are some limitations in our study that need to be addressed. First, we only used the data from the accelerometers and the timing of the cycles to characterize the biomechanical properties of adaptation in both experiments. It is possible that other aspects of the adaptation processes may be missed in this setup, such as internal foot deformations and changes in pedal and ankle rotation. We acknowledge that, in order to fully characterize the biomechanical characteristics of the adaptation process we tried to characterize at the neuromuscular level, a more comprehensive setup consisting of motion capture and force/angle-sensing pedals would be needed. Although our neuromuscular analysis suggests otherwise, we cannot exclude that a more comprehensive setup would unravel clear adaptation behaviors also for the Length experiment. The asymmetries that we tested were extremely small when compared to the biomechanics of cycling and we cannot then claim that the feedforward/feedback responses that we observed are functional. Our experimental design included a 5 minutes break between the phases of each experiment. We specifically asked subjects to remain seated and not to walk or cycle during those 5 minutes. Although previous works in literature have shown that an adapted motor plan is only forgotten if actively washed-out (e.g. (Choi and Bastian 2007)) we cannot exclude that this pause could have an effect in the adaptation/aftereffects we observed. Finally, although the results of our synergies estimation are in line with what done in similar works (Barroso et al. 2014; De Marchis et al. 2013; De Marchis et al. 2015), we acknowledge that the limited variability in the tasks we tested may affect the estimation of the muscle synergies (Steele et al. 2015).

## Acknowledgements

We thank Magdalena Gontarz for her contribution to the data collection.

## References

Barroso FO, Torricelli D, Moreno JC, Taylor J, Gomez-Soriano J, Bravo-Esteban E, Piazza S, Santos C, and Pons JL. Shared muscle synergies in human walking and cycling. J Neurophysiol 112: 1984-1998, 2014.

Berger DJ, Gentner R, Edmunds T, Pai DK, and d’Avella A. Differences in adaptation rates after virtual surgeries provide direct evidence for modularity. J Neurosci 33: 12384-12394, 2013.

Caggiano V, Leiras R, Goni-Erro H, Masini D, Bellardita C, Bouvier J, Caldeira V, Fisone G, and Kiehn O. Midbrain circuits that set locomotor speed and gait selection. Nature 553: 455–460, 2018.

Cajigas I, Koenig A, Severini G, Smith M, and Bonato P. Robot-induced perturbations of human walking reveal a selective generation of motor adaptation. Science Robotics 2: eaam7749, 2017.

Castronovo AM, De Marchis C, Schmid M, Conforto S, and Severini G. Effect of Task Failure on Intermuscular Coherence Measures in Synergistic Muscles. Applied Bionics and Biomechanics 2018: 2018.

Chia Bejarano N, Pedrocchi A, Nardone A, Schieppati M, Baccinelli W, Monticone M, Ferrigno G, and Ferrante S. Tuning of Muscle Synergies During Walking Along Rectilinear and Curvilinear Trajectories in Humans. Ann Biomed Eng 45: 1204–1218, 2017.

Choi JT, and Bastian AJ. Adaptation reveals independent control networks for human walking. Nat Neurosci 10: 1055–1062, 2007.

Chvatal SA, and Ting LH. Common muscle synergies for balance and walking. Front Comput Neurosci 7: 48, 2013.

Courtine G, Papaxanthis C, and Schieppati M. Coordinated modulation of locomotor muscle synergies constructs straight-ahead and curvilinear walking in humans. Exp Brain Res 170: 320–335, 2006.

d’Avella A, Saltiel P, and Bizzi E. Combinations of muscle synergies in the construction of a natural motor behavior. Nat Neurosci 6: 300–308, 2003.

De Marchis C, Di Somma J, Zych M, Conforto S, and Severini G. Consistent visuomotor adaptations and generalizations can be achieved through different rotations of robust motor modules. Sci Rep 8: 12657, 2018.

De Marchis C, Schmid M, Bibbo D, Castronovo AM, D’Alessio T, and Conforto S. Feedback of mechanical effectiveness induces adaptations in motor modules during cycling. Front Comput Neurosci 7: 35, 2013.

De Marchis C, Severini G, Castronovo AM, Schmid M, and Conforto S. Intermuscular coherence contributions in synergistic muscles during pedaling. Exp Brain Res 233: 1907–1919, 2015.

Dominici N, Ivanenko YP, Cappellini G, d’Avella A, Mondi V, Cicchese M, Fabiano A, Silei T, Di Paolo A, Giannini C, Poppele RE, and Lacquaniti F. Locomotor primitives in newborn babies and their development. Science 334: 997–999, 2011.

Drew T. Motor cortical cell discharge during voluntary gait modification. Brain Res 457: 181–187, 1988.

Emken JL, Benitez R, Sideris A, Bobrow JE, and Reinkensmeyer DJ. Motor adaptation as a greedy optimization of error and effort. J Neurophysiol 97: 3997–4006, 2007.

Emken JL, and Reinkensmeyer DJ. Robot-enhanced motor learning: accelerating internal model formation during locomotion by transient dynamic amplification. IEEE Trans Neural Syst Rehabil Eng 13: 33–39, 2005.

Finley JM, Bastian AJ, and Gottschall JS. Learning to be economical: the energy cost of walking tracks motor adaptation. J Physiol 591: 1081–1095, 2013.

Frigon A, Hurteau MF, Thibaudier Y, Leblond H, Telonio A, and D’Angelo G. Split-belt walking alters the relationship between locomotor phases and cycle duration across speeds in intact and chronic spinalized adult cats. J Neurosci 33: 8559–8566, 2013.

Gentner R, Edmunds T, Pai DK, and d’Avella A. Robustness of muscle synergies during visuomotor adaptation. Front Comput Neurosci 7: 120, 2013.

Grillner S. The motor infrastructure: from ion channels to neuronal networks. Nat Rev Neurosci 4: 573– 586, 2003.

Hamley EJ, and Thomas V. Physiological and postural factors in the calibration of the bicycle ergometer. J Physiol 191: 55P–56P, 1967.

Hermens HJ, Freriks B, Merletti R, Stegeman D, Blok J, Rau G, Disselhorst-Klug C, and Hägg G. European recommendations for surface electromyography. Roessingh research and development 8: 13–54, 1999.

Hug F, Turpin NA, Couturier A, and Dorel S. Consistency of muscle synergies during pedaling across different mechanical constraints. J Neurophysiol 106: 91–103, 2011.

Ivanenko YP, Cappellini G, Dominici N, Poppele RE, and Lacquaniti F. Coordination of locomotion with voluntary movements in humans. J Neurosci 25: 7238–7253, 2005.

Ivanenko YP, Poppele RE, and Lacquaniti F. Five basic muscle activation patterns account for muscle activity during human locomotion. J Physiol 556: 267–282, 2004.

Jensen L, Prokop T, and Dietz V. Adaptational effects during human split-belt walking: influence of afferent input. Exp Brain Res 118: 126–130, 1998.

Jordan LM, Liu J, Hedlund PB, Akay T, and Pearson KG. Descending command systems for the initiation of locomotion in mammals. Brain Res Rev 57: 183–191, 2008.

Kandel ER, Schwartz JH, Jessell TM, Siegelbaum SA, and Hudspeth AJ. Principles of neural science. McGraw-hill New York, 2000.

Kawato M. Internal models for motor control and trajectory planning. Curr Opin Neurobiol 9: 718–727, 1999.

Kiehn O. Decoding the organization of spinal circuits that control locomotion. Nat Rev Neurosci 17: 224–238, 2016.

Kiehn O. Locomotor circuits in the mammalian spinal cord. Annu Rev Neurosci 29: 279–306, 2006.

Lacquaniti F, Ivanenko YP, and Zago M. Patterned control of human locomotion. J Physiol 590: 2189– 2199, 2012.

Lam T, Anderschitz M, and Dietz V. Contribution of feedback and feedforward strategies to locomotor adaptations. J Neurophysiol 95: 766–773, 2006.

Lee DD, and Seung HS. Algorithms for non-negative matrix factorization. In: Advances in neural information processing systems 2001, p. 556–562.

Logan D, Ivanenko YP, Kiemel T, Cappellini G, Sylos-Labini F, Lacquaniti F, and Jeka JJ. Function dictates the phase dependence of vision during human locomotion. J Neurophysiol 112: 165–180, 2014.

MacLellan MJ, Ivanenko YP, Massaad F, Bruijn SM, Duysens J, and Lacquaniti F. Muscle activation patterns are bilaterally linked during split-belt treadmill walking in humans. J Neurophysiol 111: 1541– 1552, 2014.

Malone LA, Bastian AJ, and Torres-Oviedo G. How does the motor system correct for errors in time and space during locomotor adaptation? J Neurophysiol 108: 672–683, 2012.

McCrea DA, and Rybak IA. Organization of mammalian locomotor rhythm and pattern generation. Brain Res Rev 57: 134–146, 2008.

McDonagh MJ, and Duncan A. Interaction of pre-programmed control and natural stretch reflexes in human landing movements. J Physiol 544: 985–994, 2002.

Oliveira AS, Gizzi L, Ketabi S, Farina D, and Kersting UG. Modular Control of Treadmill vs Overground Running. PLoS One 11: e0153307, 2016.

Oliveira AS, Silva PB, Lund ME, Gizzi L, Farina D, and Kersting UG. Effects of perturbations to balance on neuromechanics of fast changes in direction during locomotion. PLoS One 8: e59029, 2013.

Pearson KG. Generating the walking gait: role of sensory feedback. Prog Brain Res 143: 123–129, 2004.

Peveler WW, Pounders JD, and Bishop PA. Effects of saddle height on anaerobic power production in cycling. J Strength Cond Res 21: 1023–1027, 2007.

Prokop T, Berger W, Zijlstra W, and Dietz V. Adaptational and learning processes during human split-belt locomotion: interaction between central mechanisms and afferent input. Exp Brain Res 106: 449– 456, 1995.

Ranganathan R, Krishnan C, Dhaher YY, and Rymer WZ. Learning new gait patterns: Exploratory muscle activity during motor learning is not predicted by motor modules. J Biomech 49: 718–725, 2016.

Reisman DS, Block HJ, and Bastian AJ. Interlimb coordination during locomotion: what can be adapted and stored? J Neurophysiol 94: 2403–2415, 2005.

Rossignol S, Dubuc R, and Gossard JP. Dynamic sensorimotor interactions in locomotion. Physiol Rev 86: 89–154, 2006.

Scheidt RA, Dingwell JB, and Mussa-Ivaldi FA. Learning to move amid uncertainty. Journal of Neurophysiology 86: 971–985, 2001.

Schillings AM, van Wezel BM, Mulder T, and Duysens J. Muscular responses and movement strategies during stumbling over obstacles. J Neurophysiol 83: 2093–2102, 2000.

Shadmehr R, and Mussa-Ivaldi FA. Adaptive representation of dynamics during learning of a motor task. J Neurosci 14: 3208–3224, 1994.

Steele KM, Tresch MC, and Perreault EJ. Consequences of biomechanically constrained tasks in the design and interpretation of synergy analyses. J Neurophysiol 113: 2102–2113, 2015.

Takakusaki K. Neurophysiology of gait: from the spinal cord to the frontal lobe. Mov Disord 28: 1483– 1491, 2013.

Torres-Oviedo G, Macpherson JM, and Ting LH. Muscle synergy organization is robust across a variety of postural perturbations. J Neurophysiol 96: 1530–1546, 2006.

Torres-Oviedo G, Vasudevan E, Malone L, and Bastian AJ. Locomotor adaptation. Prog Brain Res 191: 65–74, 2011.

Turpin NA, Guevel A, Durand S, and Hug F. Fatigue-related adaptations in muscle coordination during a cyclic exercise in humans. J Exp Biol 214: 3305–3314, 2011.

van der Linden MH, Marigold DS, Gabreels FJ, and Duysens J. Muscle reflexes and synergies triggered by an unexpected support surface height during walking. J Neurophysiol 97: 3639–3650, 2007.

Zhong G, Shevtsova NA, Rybak IA, and Harris-Warrick RM. Neuronal activity in the isolated mouse spinal cord during spontaneous deletions in fictive locomotion: insights into locomotor central pattern generator organization. J Physiol 590: 4735–4759, 2012.

